# The sensation of groove is affected by the interaction of rhythmic and harmonic complexity

**DOI:** 10.1101/415372

**Authors:** Tomas E. Matthews, Maria A. G. Witek, Ole A. Heggli, Virginia B. Penhune, Peter Vuust

**Author notes:** Corresponding author at: Department of Psychology, Concordia University 7141 Sherbrooke St W, Montreal, QC H4B 1R6.

## Abstract

Groove is defined as the pleasurable desire to move to music. Research has shown that rhythmic complexity modulates the sensation of groove but how other musical features, such as harmony, influence groove is less clear. To address this, we asked people with a range of musical experience to rate stimuli that varied in both rhythmic and harmonic complexity. Rhythm showed an inverted U-shaped relationship with ratings of pleasure and wanting to move, whereas medium and low complexity chords were rated similarly. Pleasure mediated the effect of harmony on wanting to move and high complexity chords attenuated the effect of rhythm. While rhythmic complexity is the primary driver, harmony both modulates the effect of rhythm and makes a unique contribution via its effect on pleasure. These results may be accounted for by predictive processes based on rhythmic and harmonic expectancies that are known to contribute to musical pleasure or reward.

---

When listening to music we often find ourselves tapping or moving along to the beat. This has led to the study of groove, which is the pleasurable desire to move to music[1–3]. Certain types of music are more likely to induce the sensation of groove than others. However, which specific aspects of music contribute to this sensation is less clear. Research on groove has focused on rhythmic complexity, and syncopation in particular, showing an inverted U-shaped relationship between degree of syncopation and ratings of pleasure and wanting to move [3]. That is, medium levels of syncopation are rated higher than low or high levels. This research has largely examined rhythm in isolation, but other musical properties likely contribute to the sensation of groove. Harmony in particular may be a strong contributor because it modulates affective responses [4]. However, the extent to which harmony affects groove has not been investigated. Furthermore, few studies have examined the impact of musical training on the sensation of groove. Therefore, in the current study we developed a set of rhythmic stimuli that varied in both their degree of rhythmic and harmonic complexity. We asked people with a broad range of musical training to listen to and rate how much the stimuli made them want to move and how much pleasure they experienced. Our goals were to investigate whether harmonic complexity and its interaction with rhythm affects the sensation of groove. In addition, we investigated the effects of rhythmic complexity and musical training on groove.

Early definitions of groove focused on the degree to which a piece of music will induce the desire to move to the beat [5,6]. Moving along with music, especially through dance, is often accompanied by feelings of pleasure. In a seminal study on the sensation of groove, responses to a survey emphasized both the desire to move and the associated positive affect [1]. Since then, several rhythmic aspects have been studied in terms of their effectiveness in inducing groove. Music with a strong pulse or beat leads to higher groove ratings [7,8] and is more likely to induce whole body movements, compared to music with a weak beat [9]. Percussiveness and event density have also been shown to influence the sensation of groove [7], whereas the effects of micro-timing and tempo are less clear [10–13].

A strong beat may be necessary for groove but is likely not sufficient. A ticking clock could be considered to have a strong beat but is unlikely to be something people want to dance to. Syncopation is a critical component of groove. It is often found in musical genres, such as jazz, soul, funk, Afro-Cuban, and Hip Hop [14,15], and is used by musicians intentionally to create groove [10]. An inverted U-shaped relationship has been shown between degree of syncopation, and ratings of pleasure and the desire to move, where moderately syncopated rhythms are rated as having the highest groove [3]. Syncopation occurs when a note falls on a metrically weak beat, and is then followed by a silence on a strong beat [16,17]. Meter is the pattern of strong and weak beats which may or may not be acoustically present in the rhythm. Syncopation works against this meter by emphasizing a weak beat and de-emphasizing a strong beat. This leads to violations of metric expectations [18–20] thus creating tension between the rhythm and the established meter. However, rather than reducing pleasure, listeners rate syncopated sequences as more enjoyable and sounding happier than non-syncopated sequences [21].

There are a number of non-rhythmic musical features that may contribute to the sensation of groove, including timbre, bass frequency content, and musical structure [22]. A particularly relevant potential contributor to groove is harmony because it evokes emotional valence, even for a single isolated chord [4,23,24]. Also, increasing the number of instruments, including harmony producing instruments, leads to higher groove ratings [25]. Therefore, stimuli with both rhythmic and harmonic properties are more likely to engage us, both bodily and affectively, than rhythms alone. While no current studies directly address whether harmonic complexity interacts with rhythmic complexity to affect musical enjoyment, there is evidence that consonance affects motor synchronization [26] and feelings of entrainment [27]. Therefore, in the present study we combined both rhythm and harmony to create more ecologically valid stimuli and test the effect of harmony on the sensation of groove.

To do this we varied rhythm and harmony in a similar fashion by choosing rhythms and chords that fall into three levels of complexity. Rhythms were classified based on their degree of syncopation. Chords were classified based on their degree of dissonance, which depends on both musical experience and psychoacoustic factors such as roughness and harmonicity [28–31]. Although chords most often occur in music as part of a sequence of several different chords, some groove-based genres, such as salsa, funk, and house music, frequently feature only one or two chords (James Brown’s ‘The Payback’ is a well-known example). Recent studies have shown that the harmonic complexity of isolated chords affect ratings of emotion and arousal [24] and that chords of intermediate complexity are preferred over highly consonant or dissonant chords [32]. This result supports the inverted U hypothesis which, as discussed above, has been shown for rhythmic complexity, and is theorized to be a domain-general phenomenon [33], albeit genre-dependent [34]. Additionally, isolated minor chords showed greater neural activity in emotion-related brain regions than major chords [4].

As affective, aesthetic and embodied effects of music are highly subjective and dependent on experience, musical training is likely to influence how individuals experience groove. Musicians show greater activity in emotion and reward-related brain regions while listening to music [35,36] and are more likely to employ action-based processing [37]. Further, it has been suggested that the inverted U-shaped relationship between complexity and liking disappears as musical training increases and other ‘learned aesthetic criteria’ become stronger predictors for music preference [34, pg. 608]. Several studies have suggested that musical training has little or no effect on groove ratings [3,38] or leads to lower groove ratings [25], while interest in dancing correlates positively with groove ratings [3]. Conversely, musicians have shown a greater effect of syncopation on groove ratings [39], stronger motor response to high groove music [38] and larger error-related neural response to rhythmic violations, compared to non-musicians [40]. For harmonic complexity in the context of chords, musicians show higher liking ratings [24], greater differences in ratings of consonance [4,23] and larger mismatch negativity brain responses [41]. These results suggest that musical training leads to a greater sensitivity to consonance-dissonance manipulations, but that musical training may or may not influence the effect of rhythm on groove.

Taken together, the sensation of groove involves both a motor and affective response, and is predicted by syncopation, while harmonic aspects have yet to be investigated. Harmony affects perceived emotional valence and is therefore a potential contributor to groove and may interact with rhythm. Therefore, we used an online rating study to investigate whether harmonic complexity interacts with rhythmic complexity to affect its inverted U-shaped relationship with ratings of pleasure and wanting to move. That is, to investigate whether rhythm and harmony interact to increase the subjective experience of groove in a synergistic rather than additive fashion. Effects of musical training, interest in groove music and dancing were also investigated.

## Methods

### Ethics statement

This study investigates subjective experiences of music via a web-based survey. The study was conducted through the Centre for Music in the Brain at Aarhus University, therefore, ethics were governed by the Central Denmark Region Committees on Health Research Ethics. According to their Act on Research Ethics Review of Health Research Projects (Act 593 of 14 July 2011, section 14.1), only health research studies shall be notified to the Committees. Our study is not considered a health research study (section 14.2) and therefore did not require ethical approval nor written/verbal consent, regardless of participants’ age. When recruited, participants were informed that their responses would be used for research purposes. Participants were anonymized, and no IP addresses were collected or stored. They were free to exit the survey at any time and were provided with an email address at the end of the survey to which they could address any questions or concerns.

### Participants

Two hundred and one participants between the ages of 17 and 79 (*M* = 34.74 *S*D = 13.24) completed the survey (96 reported as female). Participants reported their nationality as being from countries in six different continents, with a majority in Europe (n = 130) and North America (n = 47). As can be seen in Table S1, there was a large range of musical training backgrounds. A majority (n = 189) of participants reported no university-level music degree. Of those currently playing music, a majority played piano (n = 50), guitar (n = 44) or sang (n = 25) and had 14.5 (*SD* = 5.31) years for formal music training. Musician responders played largely classical (n = 69) or pop/rock (n = 62) genres.

Two subsets of the total sample were categorized as musicians (n = 58, 15 F) and non-musicians (n = 51, 18 F). Musicians were defined as those who reported at least eight years of formal music training (*M* = 14.5, *SD* = 5.31) and were currently practicing on a weekly or more frequent basis (hours per week: *M* = 6.52, *SD* = 8.51). Non-musicians were defined as those who reported less than three years of formal training (*M* = 0.21, *SD* = 0.49) and were not practicing on a weekly or more frequent basis.

### Stimuli

The stimuli consisted of short musical sequences that varied across three levels (Low, Medium, High) of both rhythmic and harmonic complexity. There were three different rhythms for each level of rhythmic complexity and three different chords for each level of harmonic complexity. These were combined into six versions of each rhythmic and harmonic complexity combination for a total of 54 stimuli. All stimuli were created using Cubase Pro version 8.0.30 (Steinberg Media Technologies).

Each sequence consisted of a rhythmic chord pattern with one repeated chord in a piano timbre presented at 96 beats per minute in common time (see example stimuli in Fig 1). Each sequence also included an isochronous hi-hat pattern with an inter-onset interval (IOI) of .3125 seconds, corresponding to an eighth note. The hi-hat provided a metrical context for the rhythms and prevented participants from perceptually shifting the beat of the high-complexity rhythms to reduce perceived complexity. Each piano chord lasted approximately .373 seconds including the full decay and were considered as eighth notes except in two of the high complexity rhythms which included IOI’s of .234 seconds corresponding to a dotted sixteenth note. Each sequence lasted one bar which was repeated four times for a total length of ten seconds.

**Fig 1.**
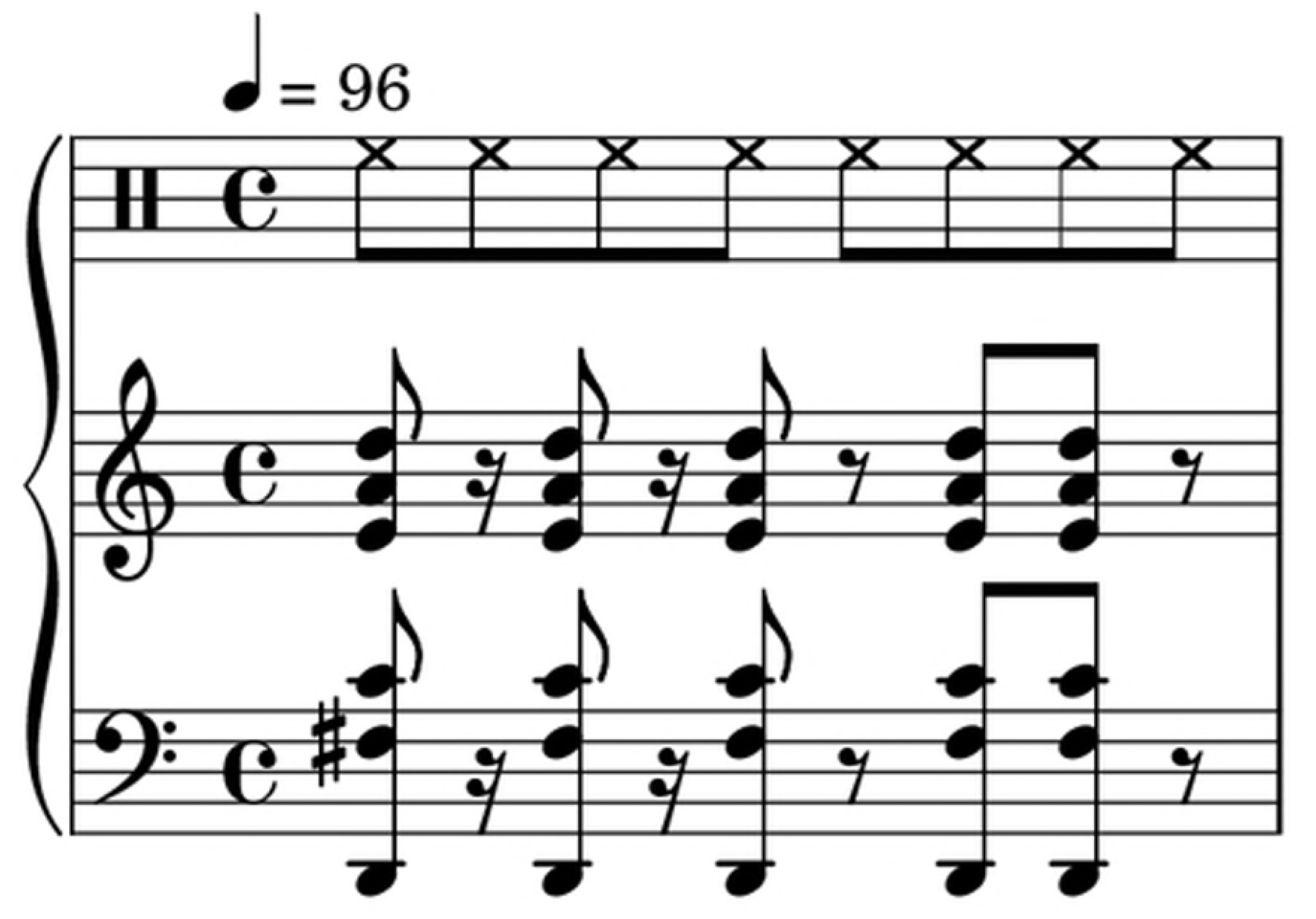
Stimuli example. Transcription of an example stimuli with a medium complexity rhythm (son clave) and a medium complexity chord (four note chord with extensions). The upper bar denotes the hi-hat.

### Rhythmic complexity

Rhythms at all three levels of complexity consisted of five onsets in a 3+2 rhythmic pattern, that is, the first half of the bar consisted of 3 onsets, and the latter of 2 onsets. Medium complexity rhythms consisted of the son clave, the rumba clave, and an experimenter-created rhythm (see Fig S1 for a schematic depiction of all rhythms). The claves were chosen as they induce a strong sense of beat despite including syncopations. The son clave and rumba clave are widely used in South American and particularly Afro-Cuban music but are also found in many forms of western music including pop, jazz and electronic dance music. Low complexity rhythms followed the same 3+2 rhythmic pattern as the medium complexity rhythms with all syncopation removed so that all onsets fall on strong beats. High complexity rhythms also followed the 3+2 rhythmic pattern, however only the first of the five onsets fell on strong beat points.

The degree of syncopation was quantified using the syncopation index created by Fitch and Rosenfeld [16] based on the formalization of syncopation by Longuet-Higgins and Lee [17]. Each syncopation in a sequence was given a weight based on the position of the rests and preceding notes involved, then these values are summed for an overall index for that sequence. The syncopation indices are summarized in Fig 2A. C-scores were also calculated for each rhythmic sequence (see Fig 2B). The C-score, created by Povel and Essens [42], is the amount of counterevidence a rhythm provides against a given metrical interpretation based on the number of weak accents and silences falling on predicted beat points. C-scores and syncopation indices were highly correlated (*r*(7) = 0.99, *p* < .05) and both were highly consistent within each level of rhythmic complexity.

**Fig 2.**
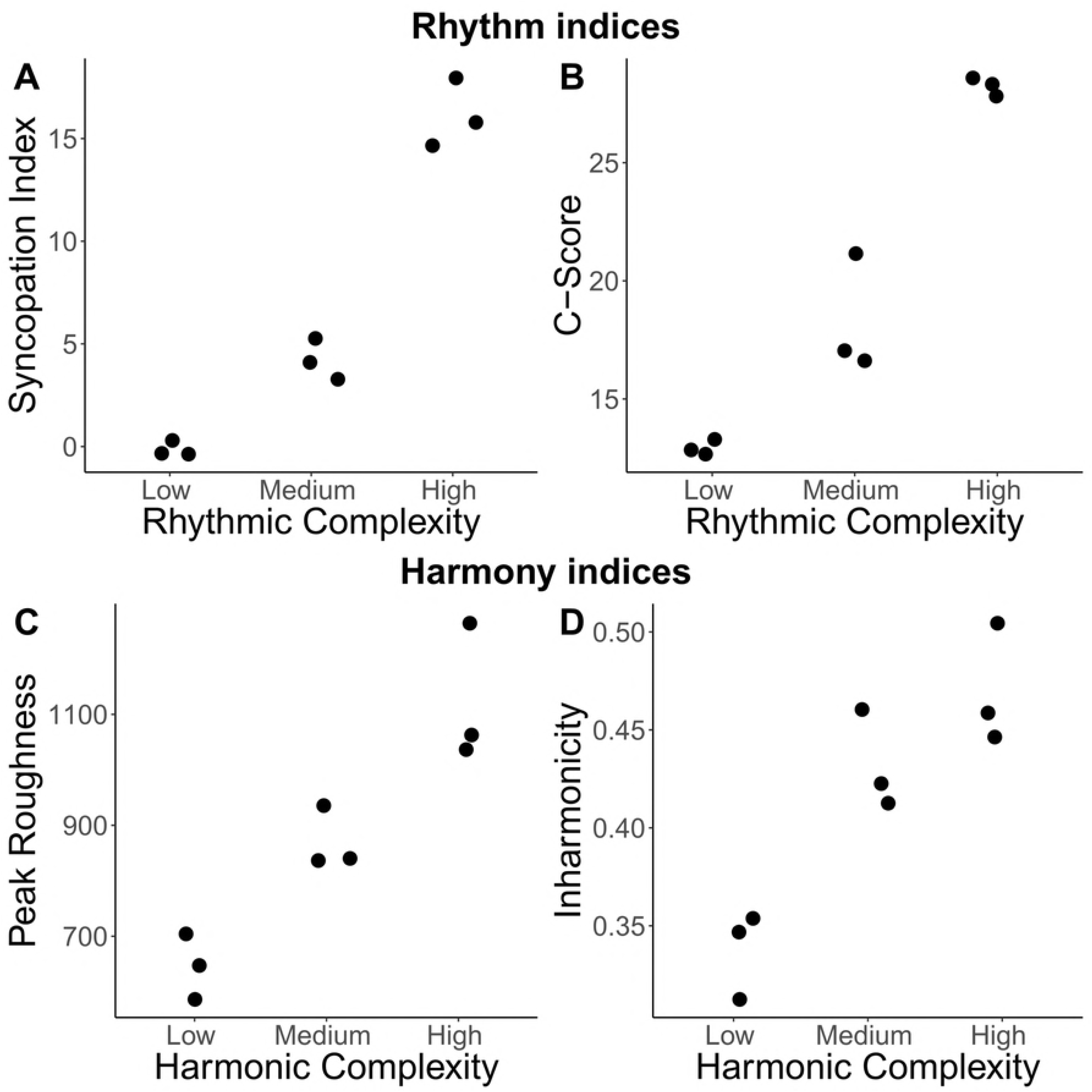
Indices of rhythmic and harmonic complexity. Scatterplots of measures of rhythmic complexity: (A) syncopation indices and (B) C-scores; and of harmonic complexity: (C) peak roughness and (D) inharmonicity.

### Harmonic complexity

There were three chords for each of the three levels of harmonic complexity (Low, Medium, and High). All chords were in the key of D major and included six notes spanning four octaves (D2 to #D5; see Fig S2). Low complexity chords consisted of the D major triad and two inversions. Medium complexity chords consisted of four note chords with extensions. High complexity chords included a flat ninth interval between chord note and extension which is considered highly dissonant, when not specifically occurring as flat 9^th^ on major 7^th^ chord, according to contemporary harmonic theory [43–45].

In order to quantify chord consonance, measures based on both acoustic and harmonic theory were used. The acoustic measures of roughness and inharmonicity were calculated with the MIRtoolbox [46]. Roughness is due to combining sounds with similar frequencies, which causes beating and sensory dissonance [47,48]. Inharmonicity refers to the degree to which the partials in a chord are integer multiples of the fundamental frequency [46]. A measure of consonance based on harmonic theory, called the aggregate dyadic consonance (ADC) [49] uses relations between pitch class sets rather than acoustic properties. Each interval class is given a consonance value which is multiplied by the number of occurrences of this interval class then summed for each chord.

As can be seen in Fig 2C, peak roughness increased with level of harmonic complexity. Mean roughness shows a similar pattern (see Fig S3). Inharmonicity increased with level of harmonic complexity, however the medium and high complexity levels showed similar values (see Fig 2D). The ADC shows an inverted U-shaped pattern where the medium complexity chords have the highest value (see Fig S3). This is related to the fact that the ADC is dependent on the number of distinct notes in a harmonic set, thus there would be more potential for consonant intervals as the number of notes increases [32,49].

## Procedure

Participants were recruited to visit a website hosting the survey via social media, email lists and word of mouth. Participants were offered the chance to win one of two Amazon gift cards worth 50 euros (or equivalent). First, participants completed a questionnaire regarding demographics, musical training, and musical preference. Participants reported how much formal training in music they had undergone, the age at which they began formal training, and how often they currently practiced. Information regarding participants’ interest in groove music, how often they listen to groove music, their enjoyment of dancing and how often they dance, were collected as ratings on a five-point scale (see Fig S4 for results). All questions required an answer before proceeding.

Participants then heard two sequences similar to the stimuli used in the survey and were asked to adjust the volume on their computer to a comfortable level. They were told to maintain the chosen volume throughout the survey. The two sequences, which were not used in the actual experiment, illustrated the range of possible levels of rhythmic and harmonic complexity. The survey then began during which each stimulus was presented once in a randomized order. After each stimulus was presented, two rating scales appeared for the two questions: ‘How much does this musical pattern make you want to move?’ and ‘How much pleasure do you experience listening to this musical pattern?’. Participants used their mouse to select their rating on the two five-point scales where one indicated ‘not at all/none’ and five indicated ‘very much/a lot’. Participants then pressed the ‘next’ button to start the next trial. Participants were not able to press ‘next’ until each stimulus had been presented in its entirety and a rating had been selected on both scales.

## Analysis

Only data from participants who completed all 54 trials were saved. Therefore, the analysis was implemented with no missing values. As ratings regarding interest and frequency of listening to groove music were highly correlated (*r*(199) = .76, 95% CI [0.696, 0.814]), they were combined using a principle component analysis (PCA) and are henceforth referred to as groove engagement PCA. The PCA loadings were then included in the models instead of the raw ratings. The identical approach was taken with the two questions regarding whether participants enjoy dancing and how often they dance (*r*(199) = .63, 95% CI [0.533, 0.703]). This variable is henceforth referred to as dance PCA.

Analysis of the main effects and interactions of rhythmic and harmonic complexity, as well as the effects of musical training, and enjoyment of dancing and groove music, were carried out using linear mixed effects regression in *R* (version 3.4.1) and *RStudio* (version 1.0.143), using the lme4 package [50]. Random intercepts for participants were included as well as by-participant random slopes for the effects of rhythm and harmony. Therefore, inter-individual differences, not only in average rating (random intercepts), but also differences in how rhythmic and harmonic complexity affected participants’ ratings (random slopes), were accounted for [51]. In addition, by-item random intercepts were included, which account for differences in ratings among the versions of stimuli within each level of rhythmic and harmonic complexity. This also allowed for analysis of the raw rather than by-level aggregated ratings. Note that Figs 3 and 5 show ratings aggregated within complexity level for visualization purposes.

A hierarchical approach was used, starting with an intercept-only model including all random effects. Predictors were then added incrementally and increases in model fit were assessed using the likelihood ratio test [52]. A final model including all significant predictors and random effects was then used to test follow up contrasts. Quadratic and pairwise contrasts were used to test whether rhythm and harmony exhibited quadratic trends and whether these trends differed across levels of the other predictors. Linear polynomial contrasts were not included as they are identical to the low versus high complexity contrast which was not of interest here. Contrasts were carried out using the emmeans package in *R* [53]. Confidence intervals were calculated using degrees of freedom approximated with the Satterthwaite method and were adjusted for multiple comparisons using the multivariate *t* method. As all contrasts involved comparing the estimates (*b*) to zero, confidence intervals not only reflect the precision of the estimate but also were used as two-tailed significance tests where an interval excluding zero indicates a significant result. Diagnostic plots of the residuals from all models were inspected for violations of the assumptions of normality and homoscedasticity. No violations were detected.

Linear regression models have been shown empirically to be robust to the potential violations of assumptions associated with Likert data [54]. However, many believe that parametric statistics such as linear mixed effects models are not appropriate for Likert data and that non-parametric and/or approaches designed for ordinal data should be used [55]. However, cumulative link mixed models (CLMM; from the ordinal package in *R*) [56], which are a standard method for analyzing ordinal data in a mixed effects context, do not allow for by-participant random slopes and are therefore less generalizable than linear mixed effect models [51]. Furthermore, simulations suggests that CLMMs are more prone to Type I errors than linear mixed effects models for Likert data [57]. In the current study, secondary analyses were carried out using CLMMs in order to compare with the linear mixed effects approach. Overall the pattern of results was very similar for both types of models with slightly more statistically significant beta estimates in the CLMM models. Given their increased generalizability and potentially lower Type I error rates compared to CLMMs, only the results of the linear mixed effects models are reported here.

### Mediation Analysis

A mediation analysis was carried out to examine whether the effects of rhythmic and harmonic complexity on *wanting to move* were mediated by *pleasure*. This involved comparing two models predicting *wanting to move* ratings. The first model included rhythmic and harmonic complexity as predictors as well as the significant covariates from the main analysis. The second, mediation model was identical to the first model, with the addition of *pleasure* ratings as a predictor. If pleasure is a significant mediator, then the contributions of rhythmic and/or harmonic complexity will be reduced in the second model. It should be noted that we did not explicitly test the directionality of the relation between pleasure and wanting to move. However, given that there is evidence that harmony in particular leads to affective responses but no evidence that harmony affects the desire to move, it seems reasonable that the relations follow the pathway from pleasure to wanting to move rather than vice versa.

The significance of the mediation effect was assessed using the mediation package [58] which provided point estimates and 95% confidence intervals for the mediation (indirect) and direct effects after taking the mediators’ effects into account. Confidence intervals were calculated using a quasi-Bayesian Monte Carlo simulation with the number of simulations set to 1000. The mediation package cannot accommodate models with maximal random effects structures therefore the models included a by-subject random intercept only. In addition, the mediation package cannot accommodate polynomial contrasts therefore only the medium versus high and medium versus low pairwise contrasts were tested.

### Group Analysis

An additional analysis was carried out to further examine the effect of musical training by comparing trained musicians with participants with little-to-no training (see Table 1 for musical background information). First, groove engagement and dance PCA loadings were compared between the musicians (n = 58) and non-musicians (n = 51). A linear mixed effects analysis tested the main effect of group, and its interactions with rhythmic and harmonic complexity, and groove engagement and dance PCA on both *wanting to move* and *pleasure* ratings.

## Results

### Wanting to move

For the *wanting to move* ratings, likelihood ratio tests that showed model fit was significantly improved by adding rhythmic complexity (*χ*^2^(2) = 280.46, *p* < .001) and harmonic complexity (*χ*^2^(2) = 134.71, *p* < .001). Follow-up contrasts showed that both rhythmic (*b*(197) = 2.269, 95% CI [-2.530, -2.008]) and harmonic complexity (*b*(199) = -0.327, 95% CI [-0.434, - 0.220]) showed significant quadratic trends, with rhythmic complexity showing a more pronounced trend. As can be seen in Fig 3A, low and medium complexity chords were rated similarly, with a drop in ratings for high complexity chords.

**Fig 3.**
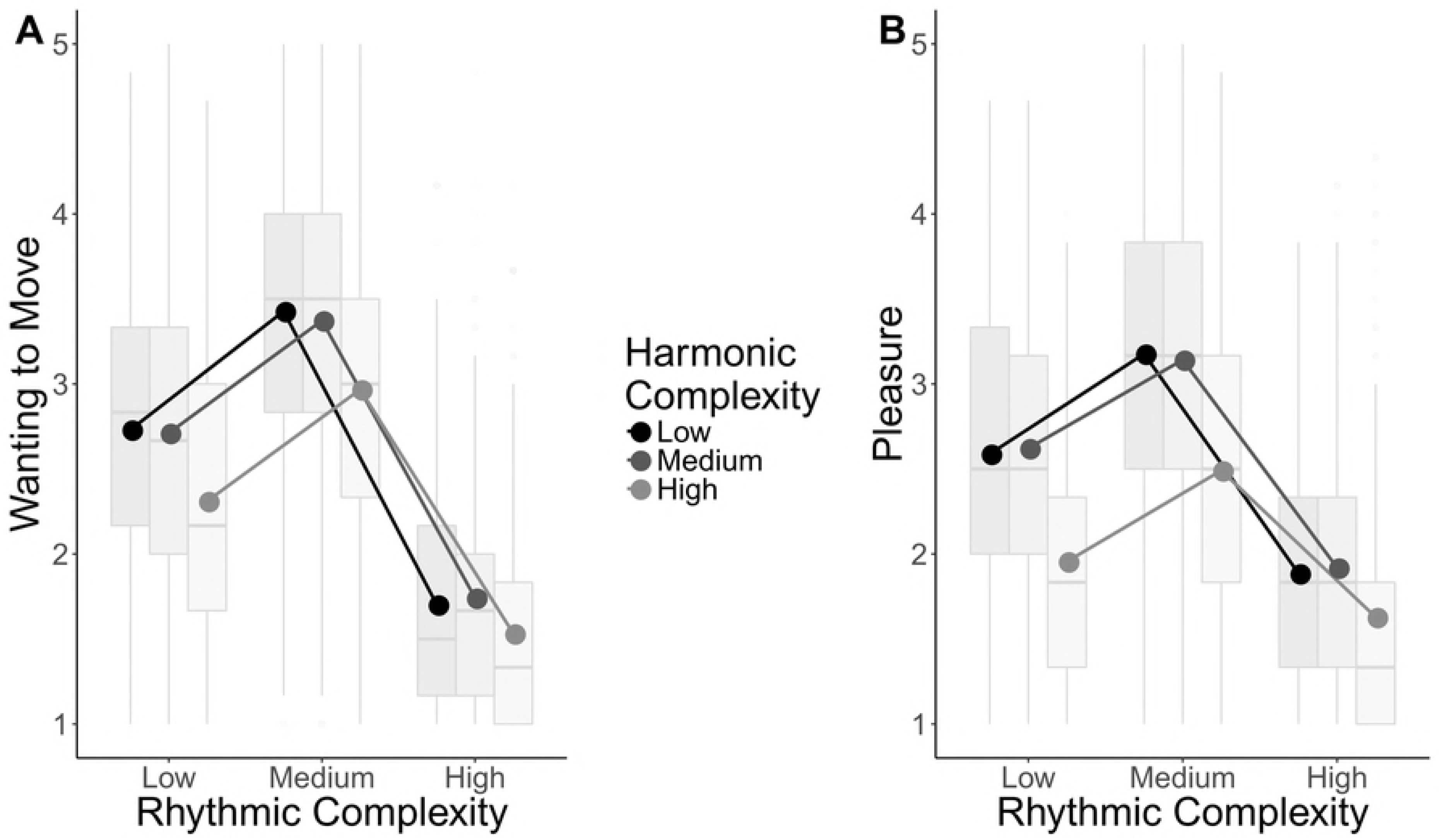
Ratings as a function of complexity. Boxplots showing the interaction between rhythmic and harmonic complexity for wanting to move ratings (A) and pleasure ratings (B). Boxplots represent ratings aggregated over items within each level of complexity for visualization purposes. Center line, median; box limits, upper and lower quartiles; whiskers, 1.5x interquartile range; points, outliers. Dots represent means calculated from the raw ratings.

Likelihood ratio tests also showed a significant interaction between rhythmic and harmonic complexity (*χ*^2^(4) = 55.27, *p* < .001). Follow-up contrasts showed that the quadratic trend for rhythmic complexity was more pronounced when combined with medium harmonic complexity than high harmonic complexity (*b*(9840) = 0.201, 95% CI [-0.008, 0.409]). However, this difference did not reach statistical significance after correction for multiple comparisons. There was a smaller difference in the quadratic trend between medium and low complexity chords which was also not significant (*b*(9840) = 0.129, 95% CI [-0.079, 0.338]). Together, these results suggest that rhythmic complexity has a more pronounced inverted U-shaped relationship with *wanting to move* when combined with low and medium complexity chords compared to high complexity chords.

Dance PCA loadings showed a significant main effect (*χ*^2^(1) = 8.30, *p* < .01). Those with greater interest in dancing showed higher *wanting to move* ratings overall (*b*(194.91) = 0.106, 95% CI [0.027, 0.186]). There were also significant interactions between dance PCA loadings and rhythmic complexity (*χ*^2^(2) = 6.97, *p* < .05), years of formal music training and rhythmic complexity (*χ*^2^(2) = 10.66, *p* < .01), and hours per week of practice and both rhythmic (*χ*^2^(2) = 8.72, *p* < .05) and harmonic complexity (*χ*^2^(2) = 10.22, *p <* .01). However, using the final model to test follow-up contrasts revealed no significant effects involving these interactions after correcting for multiple comparisons.

### Pleasure

For *pleasure* ratings, a likelihood ratio test revealed that there was a main effect of rhythmic complexity (*χ*^2^(2) = 227.49, *p* < .001). When harmonic complexity was added, the model failed to converge (Barr et al., 2013). Therefore, the main effect of harmonic complexity was added at the same step as the harmony by rhythm interaction, which together significantly improved model fit (*χ*^2^(6) = 295.69, *p* < .001). Follow-up contrasts showed that both rhythmic (*b*(197) = -1.673, 95% CI [-1.905, -1.442]) and harmonic complexity (*b*(198) = -0.546, 95% CI [-0.688, -0.405]) showed significant quadratic trends. As in the *wanting to move* results, rhythm showed a pronounced inverted U shape, whereas low and medium complexity chords were rated similarly and with a drop in ratings for high complexity chords (see Fig 3B).

Follow-up contrasts for the rhythm by harmony interaction showed that the quadratic trend for rhythmic complexity was significantly more pronounced for medium than high complexity chords (*b*(9840) = 0.345, 95% CI [0.131, 0.559]). There was a smaller, non-significant difference in the trend between medium and low complexity chords (*b*(9840) = 0.138, 95% CI [-0.075, 0.352]). Therefore, like the *wanting to move* results but more prominent, the inverted U relationship between rhythm complexity and *pleasure* was more pronounced for low and medium complexity chords compared to high complexity chords.

Likelihood ratio test showed significant interactions between hours of weekly practice and both rhythmic (*χ*^2^(2) = 14.63, *p* < .001) and harmonic complexity (*χ*^2^(2) = 12.04, *p* < .01). There was a significant interaction between dance PCA loadings and rhythmic complexity (*χ*^2^(2) = 7.16, *p* < .05). When adding groove engagement PCA loadings, the model failed to converge, therefore, groove engagement PCA loadings were added at the same step as the rhythm by groove engagement PCA interaction, which increased model fit (*χ*^2^(3) = 8.81, *p* = .03). There was also a significant interaction between groove engagement PCA loadings and harmonic complexity (*χ*^2^(2) = 6.73, *p* < .05). However, using the final model to test follow-up contrasts revealed no significant effects involving these main effects or interactions after correcting for multiple comparisons.

### Mediation Analysis

Two models predicting *wanting to move* ratings were used to test whether pleasure mediated the effect of harmonic and rhythmic complexity on wanting to move. The first model included all the significant covariates and interactions from the main analysis (rhythm, harmony, Dance PCA, years of music training and hours per week of practice), and the second, mediation model was identical to the first with the addition of pleasure ratings as a predictor.

For rhythmic complexity, *pleasure* ratings had a significant mediation effect while the direct effect of rhythmic complexity was also significant. Both effects are evident in that the difference in *wanting to move* ratings between medium and low rhythmic complexity was significant in both the first model (*b*(1592) = 0.672, 95% CI [0.585, 0.759]) and reduced, but still significant in the mediation model (*b*(1634.9) = 0.294, 95% CI [0.227, 0.361]) showing the remaining direct effect. The reduction of the effect from the first to the mediation model was itself significant (*b* = 0.378, 95% CI [0.329, 0.430]), suggesting *pleasure* had a mediating effect. The difference in ratings between the medium and high complexity rhythms showed the same pattern. The difference was smaller in the mediation model (*b*(1729.54) = 0.821, 95% CI [0.741, 0.900]) compared to the initial model (*b*(1592) = 1.60, 95% CI [1.510, 1.684]), and this decrease was significant (*b* = 0.777, 95% CI [0.718, 0.840]). Therefore, for both the medium versus low and medium versus high rhythm complexity contrasts, *pleasure* showed a significant mediation effect, while the direct effect remained significant.

For harmonic complexity, the difference in ratings between medium and low complexity chords was not significant in the initial model (*b*(1592) = 0.012, 95% CI [-0.075, 0.099]) or the mediation model (*b*(1591.03) = 0.019, 95% CI [-0.044, 0.081]). For the medium minus high harmonic complexity contrast, *pleasure* was a significant mediator as the contrast was significant in the first model (*b*(1592) = 0.339, 95% CI [0.252, 0.426]) and not in the mediation model (*b*(1633.31) = -0.031, 95% CI [-0.098, 0.036]), therefore no direct effect remained. The mediating effect of *pleasure* ratings was significant (*b* = 0.371, 95% CI [0.323, 0.420]).

These results, summarized in Fig 4, show that *pleasure* ratings fully mediated the effect of harmonic complexity on *wanting to move* ratings. However, *pleasure* only partially mediated the effect of rhythmic complexity on *wanting to move* ratings such that a direct effect of rhythmic complexity remained.

**Fig 4.**
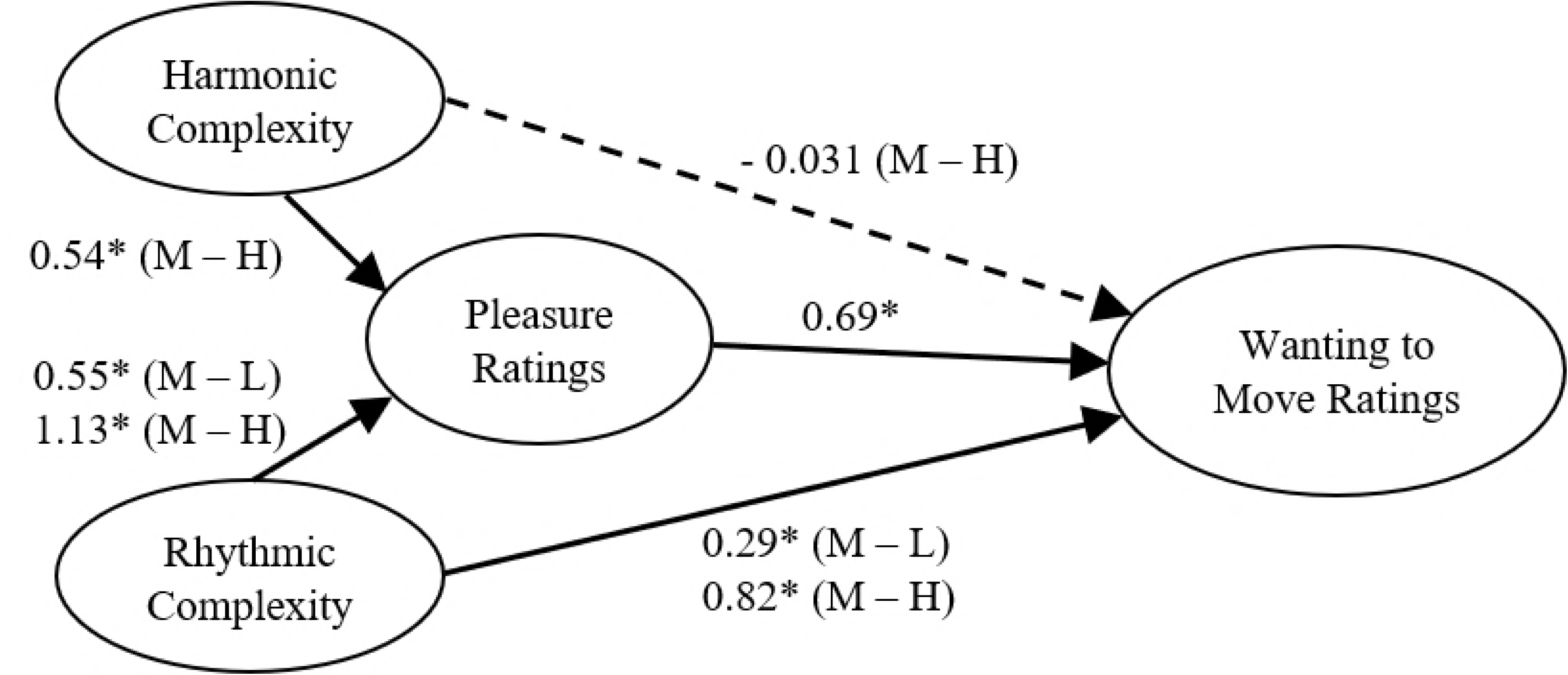
Path model. Path model based on the mediation analysis showing the relations between the predictors – rhythmic and harmonic complexity; the mediator – pleasure ratings; and the outcome variable – wanting to move ratings. Regression estimates for the effects of rhythmic and harmonic complexity on wanting to move ratings are from the mediation model that takes into account the effect of pleasure ratings on wanting to move ratings. The dashed line indicates that the direct effect of the medium – high harmonic complexity contrast was no longer significant once pleasure ratings were included in the model. L = Low, M = Medium, H = High; * *p* < .05.

### Musician vs non-musicians

Dance PCA loadings were not significantly different between groups (*b*(106.31) = 0.324, 95% CI [-0.257, 0.906]), nor were groove engagement PCA loadings (*b*(105.61) = - 0.099, 95% CI [-0.675, 0.477]).

***Wanting to move***

There was no significant main effect of group (*χ*^2^(1) = 0.25, *p* > .05), but there was a significant interaction between group and rhythmic complexity (*χ*^2^(2) = 7.47, *p* < .05) on *wanting to move* ratings. A follow-up contrast showed that the quadratic trend for rhythmic complexity was more pronounced for musicians than non-musicians (*b*(103) = 0.780, 95% CI [0.105, 1.456]; see Fig 5A). There was no significant three-way interaction between group, rhythmic and harmonic complexity. Although there were significant three-way interactions between group, rhythmic complexity and both dance PCA loadings (*χ*^2^(4) = 11.14, *p* < .05) and groove engagement PCA loadings (*χ*^2^(4) = 11.03, *p* < .05), follow up contrasts corrected for multiple comparisons revealed no main effects or interactions.

#### Pleasure

There was a significant effect of group (*χ*^2^(1) = 4.52, *p* < .05) showing that musicians had higher *pleasure* ratings overall compared to non-musicians (*b*(107) = 0.227, 95% CI [0.023, 0.431]; see Fig 5B).

**Fig 5.**
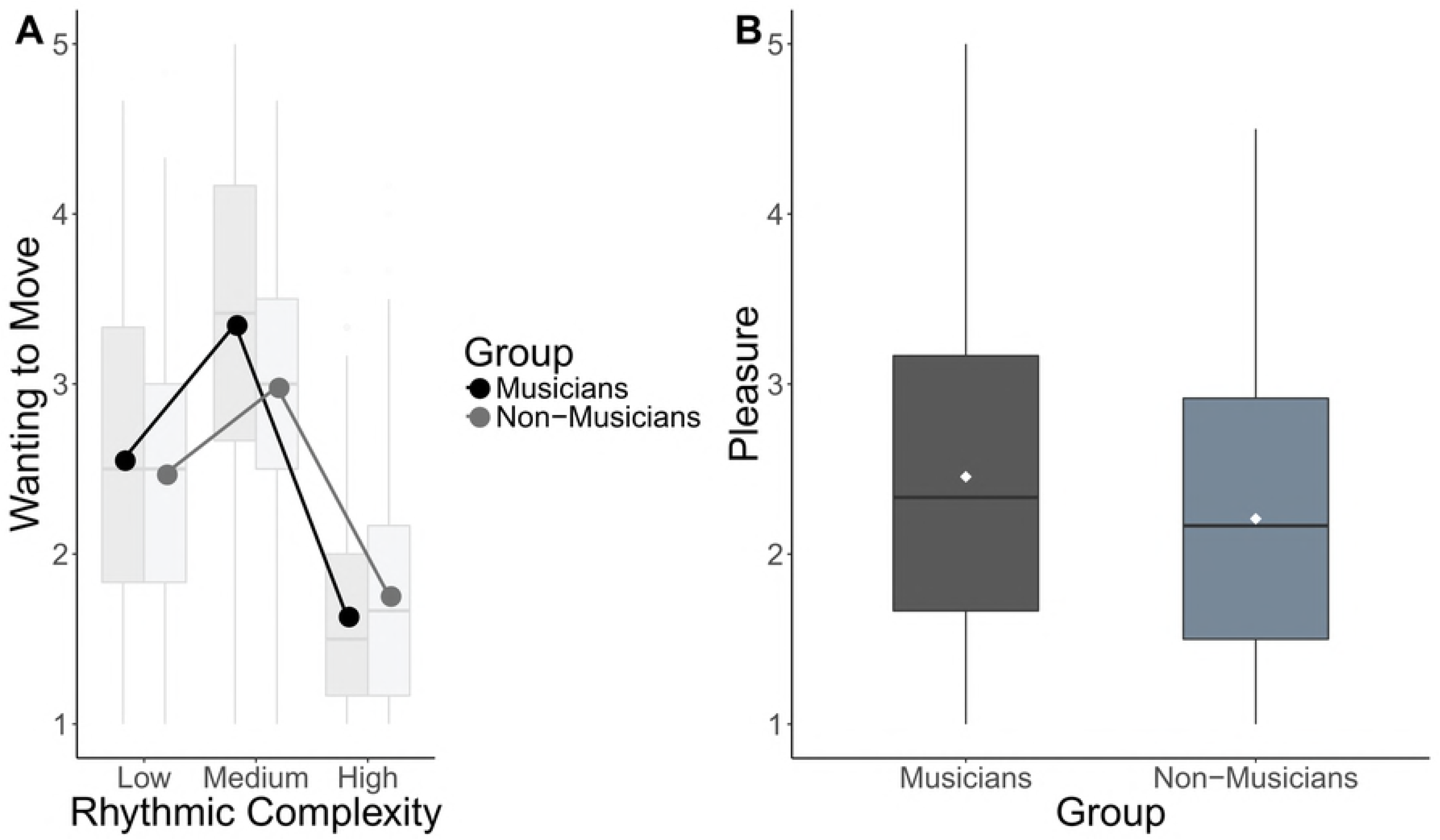
Ratings as a function of musical training. A) Box plot showing the interaction between group and rhythmic complexity. Lines represent means calculated from raw ratings. B) Box plot the effect of musical training on pleasure ratings. Boxplots represent ratings aggregated over items within each level of complexity for visualization purposes. Center line, median; box limits, upper and lower quartiles; whiskers, 1.5x interquartile range; points, outliers. Dots represent means calculated from the raw ratings.

## Discussion

This study used ratings of *pleasure* and *wanting to move* to assess whether harmony and rhythm work together to affect the sensation of groove. Rhythm showed a strong inverted U-shaped relationship with both pleasure and wanting to move ratings. Harmony did not show an inverted U-shaped relationship as medium and low complexity chords were rated similarly. Rhythm and harmony interacted such that the inverted U-shaped pattern for rhythm was less prominent in the context of high compared to medium and low complexity chords, particularly for *pleasure* ratings. That is, high harmonic complexity not only reduced the sensation of groove overall but reduced the effect of rhythm complexity as well. Furthermore, the effect of harmony on wanting to move was strongly mediated by pleasure, while rhythm directly affected both wanting to move and pleasure. Together these results suggest that rhythm plays a primary role in generating the sensation of groove, with harmony providing a modulatory role through its effect on pleasure.

Importantly, musicians showed a stronger effect of rhythmic complexity on *wanting to move* ratings and higher *pleasure* ratings overall. Together these results show that musical training strengthens the connection between syncopation and the desire to move and leads to greater reported pleasure. Finally, for all participants, interest in dance was associated with higher *wanting to move* ratings.

Consistent with the previous literature, rhythmic complexity showed an inverted U-shaped relationship with the sensation of groove such that medium levels of syncopation increased pleasure and desire to move compared to low and high levels [3,59]. Emotional and embodied responses to music have long been thought to result from predictive processes whereby listeners develop internal models, or musical expectancies, based on prior experience [18–20]. The strongest responses arise when listeners can make predictions, but expectancies are subtly violated, creating a balance between predictability and uncertainty [60]. According to the predictive coding framework, medium levels of syncopation achieve this balance by creating an optimal level of tension between a predictive model – the meter – and the current sensory input – the rhythm [40,61,62]. If the rhythm is too complex, no model of the meter can be established and if it is too simple, there is no tension. This tension has been hypothesized to increase pleasure because it engenders prediction errors, or violations of expectations, which are rewarding as they lead to further predictions and thus learning [62]. The desire to move is highest for medium syncopation because the tension between model and input encourages the listener to reinforce this model by synchronizing their movements which fills in the gaps in the rhythmic surface created by syncopations [63]. Movement may also be a way for listeners to test their model of the meter [64].

While it is not yet clear how pleasure and desire to move interact to create the sensation of groove, theories of entrainment offer promising hypotheses. Motor brain regions are important for beat perception [65–68] and entrainment of activity in motor and auditory regions to rhythmic stimuli drive auditory temporal predictions [69], as well as meter and beat perception [70–73]. Motor regions may also be involved in the affective response to music as they show increased activity for preferred over non-preferred rhythms [74] and during music-induced affective responses [75]. Finally, motor cortical excitability has been shown to increase when people listened to high groove music [38]. More generally, feelings of entrainment also predict positive affective responses to music [76]. This is in line with the idea that entrainment at neural, cognitive, physiological, and social levels results in a positive affective response [77]. In the current study, medium complexity rhythms may have engendered greater entrainment at one or more of these levels resulting in increased pleasure and desire to move. It is also possible that familiarity may have contributed to the U-shaped relationship observed here because the medium complexity rhythms consist of son and rumba claves that are common to many types of popular music. However, the stimuli used here were entirely novel, and therefore would not be individually recognizable.

Compared to rhythm complexity, harmonic complexity showed a less marked U-shaped pattern for both wanting to move and pleasure, because low and medium complexity chords were rated similarly. This may be because low and medium complexity chords are both relatively common in groove music, and thus did not differ in pleasure. In contrast, high complexity chords are uncommon, and thus were not only perceived as unpleasant but violated expectations, resulting in a strong negative effect on both pleasure and the desire to move. In addition, based on our findings, rhythmic features appear to dominate for these stimuli, which may have reduced the attention paid to harmonic complexity. Another possibility is that the range of harmonic complexity was not large enough to capture an inverted U-shaped relationship. Lower complexity chords, in this case the octave, may lead to lower ratings than the low complexity chords used here.

High complexity chords attenuated the effect of rhythm complexity on the sensation of groove. This attenuation was significant for pleasure and only near-significant for wanting to move ratings which, combined with the results of the mediation analysis, suggests that harmony primarily affects the pleasure component of groove. These effects may be due to a shared internal model that generates predictions about both the timing and harmonic content of future events. Behavioural evidence for a shared model is found in a study showing more accurate pitch judgements for rhythmically expected tones [78]. In addition, neural beta oscillations entrained to a rhythmic stimulus are sensitive to violations of both rhythmic [79] and pitch expectations [80]. Shared internal models enhance predictive processing [69], however violations of one component of the model likely affect the other. As discussed above, high complexity chords violate harmonic expectations, which may disrupt the shared model and increase the disparity between model and sensory input, thus reducing pleasure and the desire to move.

As discussed above, pleasurable states are thought to facilitate entrainment at various levels [77]. According to this view, harmonic complexity may affect the degree of entrainment at one or more of these levels via its influence on pleasure. This is in line with the result that pleasure ratings strongly mediated the effect of harmonic complexity on wanting to move ratings. This suggests that harmonic complexity does not directly influence the desire to move and instead affects the experienced pleasure, which in turn affects the desire to move. The effect of harmony on pleasure may therefore influence the degree of entrainment, thus affecting the desire to move while also modulating the effect of rhythm complexity. Support for this hypothesis comes from work showing that auditory-motor synchronization was reduced for dissonant compared to consonant tones [26] and that consonant music generated greater feelings of entrainment [27]. Conversely, feelings of entrainment have also been found to enhance pleasure [76]. We hypothesize that the desire to move and pleasure associated with groove are reciprocal and based on interactions between predictive and entrainment processes in the auditory, motor, and reward systems.

Compared to non-musicians, musicians showed a more prominent inverted U-shaped relationship between rhythm and wanting to move. Musical training may lead to an increased awareness and appreciation of syncopation and its effect on the desire to move. For example, musicians have been shown to use syncopation intentionally to convey groove [10] and musical expertise has been positively linked with the effect of syncopation on groove ratings [39]. In addition, musical training may lead to more developed internal models that lead to stronger rhythmic expectations. This is supported by studies showing that musicians have greater error-related neural responses to rhythmic violations [40] and enhanced neural entrainment to natural music [81]. An increase in motor-based processing may also account for the greater effect of rhythm on wanting to move in musicians [37,38]. Consistent with previous work [3], enjoyment and interest in dancing was also associated with higher wanting to move ratings overall. This further supports the link between motor processes and groove-based music in those with strong associations between music and movement.

The musician group also showed greater overall pleasure ratings compared to non-musicians. This is consistent with evidence that musicians demonstrate greater enjoyment of and increased neural reward activity for a range of musical stimuli [24,35,36]. Some studies have shown no effect of musicianship [3], or reduced groove ratings in musicians [25]. However, these studies defined musicianship less strictly, thus perhaps attenuating the effects of training-based internal models or expectancies on the sensation of groove.

We have shown that rhythm and harmony interact in the sensation of groove. While rhythmic complexity is the primary driver, harmony both modulates the effect of rhythm and makes a unique contribution via its effect on pleasure. These results can be accounted for by predictive processes based on rhythmic and harmonic expectancies. Syncopated rhythms create the optimal level of tension between expectancy and violation which increases pleasure and the desire to move. Harmonic expectancies also affect pleasure, and by influencing emotional valence, enhance the effects of rhythm. Musical expectancies are encoded in auditory-motor networks and influenced by experience, which may account for the increased sensitivity to groove in those with strong associations between movement and music. Taken together, this work provides important new information about the role of prediction in the experience of musical pleasure. These findings may also contribute to the development of more effective and enjoyable music-based interventions.

## Data Availability

The ratings and background data that support these findings are available in the Open Science Framework with identifier link: DOI 10.17605/OSF.IO/76ZWY

## Supporting Information

**Table S1. Musical Background.** (DOCX)

**Fig S1. Bar plots of demographic information.** (A) Number of participants in from each continent; AF, Africa; AS, Asia; EU, Europe; NA, North America; OC, Oceania; SA, South America. (B) Number of participants with each type of degree in music; Bach, Bachelors. (C) Type of instrument played musicians; DJ/Prod, DJ/Producer. (D) Genre played by musicians; D/E, Dance/Electronic; Exp, Experimental; N-W, non-western. (DOCX)

**Fig S2**. **Bar plots of groove and dance questions.** Questions regarding groove (A and B); Questions regarding music and dancing (C and D). (DOCX)

**Fig S3. Schematic representation of rhythms used to create the stimuli.** Weights represent weights used to calculate the syncopation index. Medium 1 = Son clave, Medium 2 = Rumba clave. (DOCX)

**Fig S4. Chords used in the stimuli**. a) low harmonic complexity, b) medium complexity chords, c) high complexity chords. DOCX

**Fig S5. Additional indices of harmonic complexity.** (A) Mean roughness, and (B) ADC indices for the chords used in the stimuli.

## Acknowledgements

The authors thank Alessandra Brusa for help with the dissemination of the survey. They also thank Gabe Nespoli for analysis suggestions and Elvira Brattico for practical help and advice.

## Author Contributions

Conceptualization: TM, MW, PV, VP. Data Curation: OH, TM, MW. Formal Analysis: TM. Funding Acquisition: TM, PV, VP. Investigation: TM, MW, OH. Methodology: TM, MW. Project Administration: TM, MW. Resources: OH, PV. Software: OH, TM, MW. Supervision: TM, MW. Validation: TM. Visualization: TM. Writing – Original Draft Preparation: TM. Writing – Review & Editing: TM, MW, PV, VP.

